# Prevalent co-release of glutamate and GABA throughout the mouse brain

**DOI:** 10.1101/2024.03.27.587069

**Authors:** Cesar C. Ceballos, Lei Ma, Maozhen Qin, Haining Zhong

**Affiliations:** Vollum Institute, Oregon Health & Science University, Portland, OR 97239, USA

## Abstract

Several neuronal populations in the brain transmit both the excitatory and inhibitory neurotransmitters, glutamate, and GABA, to downstream neurons. However, it remains largely unknown whether these opposing neurotransmitters are co-released onto the same postsynaptic neuron simultaneously or are independently transmitted at different time and locations (called co-transmission). Here, using whole-cell patch-clamp recording on acute mouse brain slices, we observed biphasic miniature postsynaptic currents, i.e., minis with time-locked excitatory and inhibitory currents, in striatal spiny projection neurons (SPNs). This observation cannot be explained by accidental coincidence of monophasic miniature excitatory and inhibitory postsynaptic currents (mEPSCs and mIPSCs, respectively), arguing for the co-release of glutamate and GABA. Interestingly, these biphasic minis could either be an mEPSC leading an mIPSC or vice versa. Although dopaminergic axons release both glutamate and GABA in the striatum, deletion of dopamine neurons did not eliminate biphasic minis, indicating that the co-release originates from another neuronal type. Importantly, we found that both types of biphasic minis were detected in other neuronal subtypes in the striatum as well as in nine out of ten additionally tested brain regions. Our results suggest that co-release of glutamate and GABA is a prevalent mode of neurotransmission in the brain.

## INTRODUCTION

The long-accepted Dale’s principle postulated that each neuron releases a single transmitter type (1, 2). However, recent works have demonstrated that certain neuronal populations are capable of releasing two or more neurotransmitters (3–5). There are two qualitatively distinct modes of releasing multiple neurotransmitters from the same neurons. They can be simply “co-transmitted”, i.e., two transmitters being released from different vesicles at different locations and/or times. Alternatively, they can be “co-released” simultaneously at the same location, typically from the same vesicle. These two modes of release have distinct implications for how the two neurotransmitters coordinate each other for function (3, 4).

Among the multimodal neurons, a peculiar subset can co-transmit glutamate and GABA, the primary excitatory and inhibitory neurotransmitters in the brain, respectively (6–16). How these two neurotransmitters are released are being intensively studied. Given the opposing functions of these two neurotransmitters, it would seem logical if they are released from different synaptic pools. However, recent evidence at the lateral habenula suggest that glutamate and GABA can co-released from the same synaptic vesicles (9, 17). Furthermore, single-cell RNA sequencing and immunostaining studies suggest that vesicular transporters for both glutamate and GABA (VGAT and VGLUT1/2/3, respectively) are co-expressed in many neuronal populations throughout the brain, sometimes even on the same synaptic vesicles (18–24). However, it remains unclear whether co-release is a general mechanism for glutamate/GABA bimodal neurons and whether such co-release occurs broadly in the brain.

We set out to answer these questions first in the dorsolateral striatum. It is well established that dopaminergic axon terminals from Substantia Nigra pars compacta (SNc) can release glutamate and GABA in addition to dopamine in the striatum (7, 25–29), but whether these neurotransmitters are co-released is still under investigations (11, 30–35). We recorded miniature postsynaptic currents (minis), which are thought to arise from the release of neurotransmitters by a single vesicle. At a holding voltage that can detect both excitatory and inhibitory synaptic currents, we observed “biphasic” minis in striatal projection neurons (SPNs) that depended on both AMPA and GABA_A_ receptors and occurred beyond the co-incidence chance expected of independent excitatory and inhibitory minis. The frequency of biphasic minis could also be modulated independently from those of monophasic minis. Minimal optogenetic stimulation experiments and neuronal ablation experiments demonstrated that these biphasic minis did not depend on the mid-brain dopaminergic neurons. Surprisingly, such biphasic minis were detected in all but one of 12 brain regions or cell types that we have examined. Our results suggest that glutamate and GABA co-release is a prevalent feature of a subset of synapses throughout the brain.

## RESULTS

### Detection of biphasic minis in SPNs of the dorsolateral striatum

Biphasic miniature synaptic currents, mediated by both AMPA and GABA_A_ receptors may indicate co-release from the same synaptic vesicles (9, 36–38). To determine whether such co-release occurs in the dorsolateral striatum, we recorded minis from putative SPNs in acute brain slices of wildtype mice using whole-cell patch-clamp recording in the presence of TTX (1 µM). Both miniature excitatory and inhibitory postsynaptic currents (mEPSCs and mIPSCs, respectively) were observed in the same trace when the cell was held at -30 mV, but less so at -10 or -50 mV (Fig. 1a, 1b and Supplementary Fig. 1). Careful examination of the traces at -30 mV revealed a unique type of minis that exhibited a clear biphasic shape (Fig. 1c–1f). Near all of these minis (181 out of 184) were well fit by an averaged mEPSC followed by an mIPSC with a small delay (Fig. 1g; delay = 9.1 ± 3.9 ms, mean ± s.d.). These biphasic minis were blocked by either the AMPA receptor (AMPAR) antagonist NBQX (10 µM) or the GABA_A_ receptor (GABA_A_R) antagonist GABAzine (10 µM) (Fig. 1d and 1e). As the holding voltage was moved from -50 to -10 mV, the inward component decreased and the outward component increased (Fig. 1h and 1i). These results indicate that the biphasic minis are the result of near simultaneous activation of AMPA and GABA_A_ receptors.

**Fig. 1.**
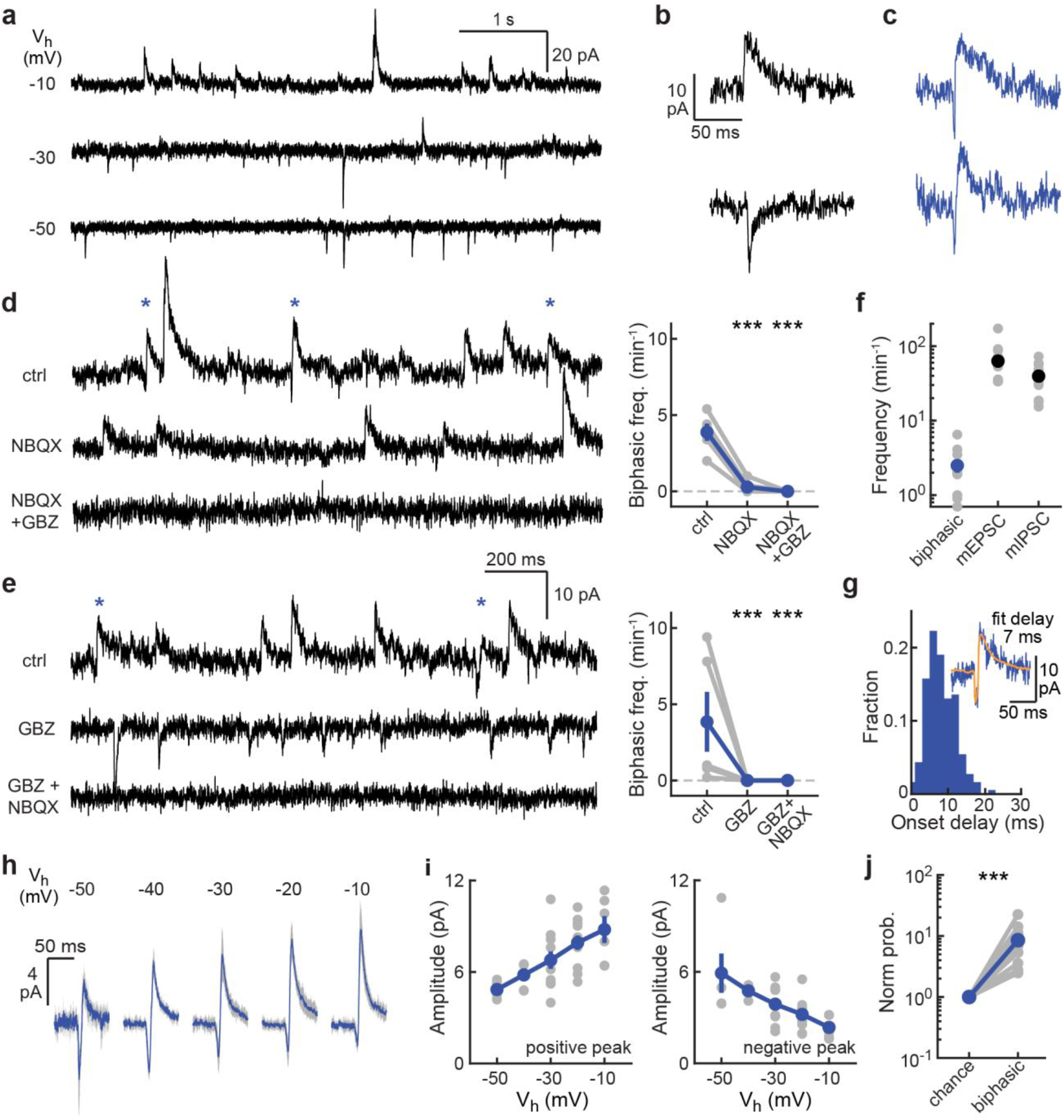
Detection of straight biphasic minis in SPNs of the dorsolateral striatum. **a**, Example traces of miniature current recordings at different holding potentials (V_H_). **b**, Example traces of mIPSC (top) and mEPSC (bottom). **c**, Example traces of straight biphasic minis at V_H_ = -30 mV. **d** and **e**, Biphasic minis are abolished by either AMPAR (**d**) or GABA_A_R (**e**) antagonists (NBQX or GABAzine, respectively). n (neurons/slices/mice) = 5/5/3 for both. **f**, Frequency (# events/min) of biphasic minis, mEPSCs and mIPSCs at V_H_ = -30 mV. **g**, Histogram of the onset delay between the mEPSC and mIPSC component used to fit the biphasic minis. n = 181 events. Mean ± s.d. = 9.1 ± 3.9 ms. Inset, biphasic mini fitted by a 7-ms delay. **h**, Average traces of biphasic minis at different holding potentials. **i**. Voltage-dependent changes of the amplitude of the maximum and minimum peaks of the straight biphasic minis. **j**, The probability of observed straight biphasic minis normalized to the probability of their occurrence due to coincidence of independent mEPSC and mIPSC. n (neurons/slices/mice) = 11/7/4 for panels **h**–**j**.

To test whether biphasic minis were caused by coincided arrival of independent monophasic mEPSCs and mIPSCs, we calculated the likelihood of mEPSCs and mIPSCs arriving in the same time window (10 ms; see Fig. 1g) using the observed mEPSC and mIPSC frequencies (see Methods), and compared to the probability of biphasic minis from our recordings. The frequency of biphasic minis was significantly lower than either mEPSCs or mIPSCs (Fig. 1f), but was still about 10-fold higher than chance (p < 0.001; Fig. 1j), suggesting that biphasic minis are the result of correlated release of glutamate and GABA.

Interestingly, in the same recording, we also observed biphasic minis in which the outward current preceded the inward current (Fig. 2a and 2b). We called this second type “reverse” biphasic minis, whereas the original type as “straight”. The reverse biphasic minis were also sensitive to the blockade of either AMPARs or GABA_A_Rs (Fig. 2c and 2d), and were well fit by an mIPSC preceding a mEPSC, with an average delay of 11.1 ± 3.9 ms (mean ± s.d.; Fig. 2e). Reverse biphasic minis also occurred well above the chance of coincided independent mEPSCs and mIPSCs (p < 0.01; Fig. 2f), suggesting that they reflected correlated release of glutamate and GABA.

**Fig. 2.**
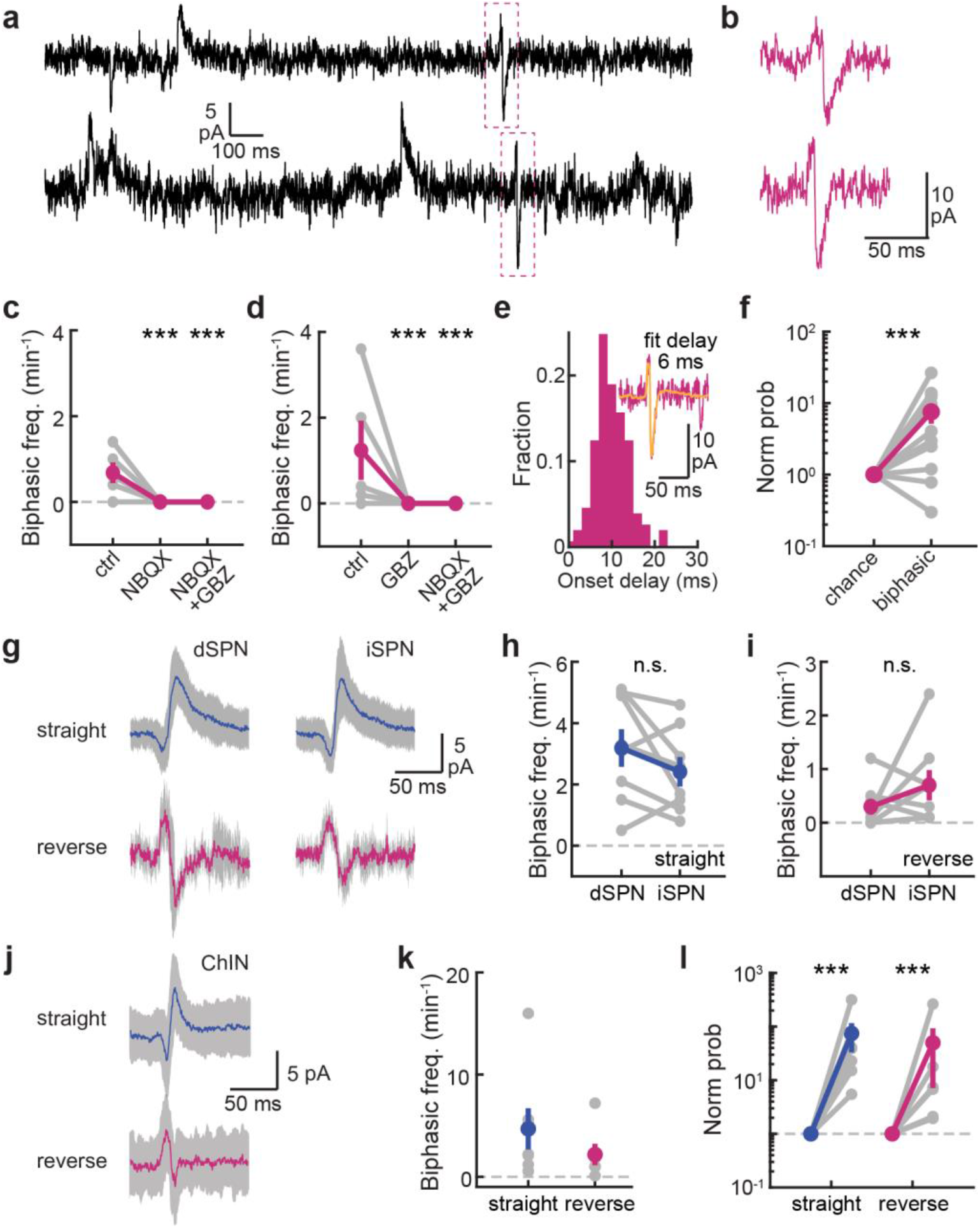
Reverse biphasic minis in SPN of the dorsolateral striatum. **a**, Example traces with reverse biphasic minis (dashed red box). **b**, Example traces of reverse biphasic minis at V_H_ = -30 mV. **c** and **d**, Reverse biphasic minis are abolished by NBQX (**c**) or GABAzine (GBZ; **d**). n (neurons/slices/mice) = 5/5/3 for both. **e**, Histogram of the onset delay between the mEPSC and mIPSC component used to fit the biphasic minis. N = 151 events. Mean ± s.d. = 11.1 ± 3.9 ms. Inset, a reverse biphasic mini fitted by a 6-ms delay. **f**, The probability of observed reverse biphasic minis normalized to the probability of their occurrence due to coincidence of independent mEPSC and mIPSC. n (neurons/slices/mice) = 11/7/4. **g**, Example average traces of straight and reverse biphasic minis recorded in dSPNs and putative iSPNs. **h** and **i**, Frequency (events/min) of straight (**h**) and reverse biphasic minis (**i**) from paired dSPNs and iSPNs at V_H_ = -30 mV. n (neurons/slices/mice) = 8/5/2. **j**–**l**, Example average traces (**j**), frequency (**k**), and occurrence probability normalized to chance (**l**) of straight and reverse biphasic minis recorded in ChIN at V_H_ = -30 mV. n (neurons/slices/mice) = 6/3/2.

To examine whether biphasic minis may be specific to a unique neuronal type in the striatum. we used the *D1R-tdTomato* mouse line to distinguish direct pathway SPNs (dSPNs) from putative indirect-pathway SPNs (iSPNs; i.e., tdTomato-negative neurons that were not cholinergic interneurons [ChINs], which have a unique large soma). Both types of biphasic minis were present in dSPNs and iSPNs at comparable frequencies that were above chance (Fig. 2g–2i). Furthermore, both straight and reverse biphasic minis were also present in ChINs (Fig. 2j–2l). Thus, biphasic minis of both types appeared to be broadly present in multiple neuronal types in the striatum.

### Biphasic minis are independently modulated compared to monophasic minis

To further determine whether the biphasic minis are independent from monophasic mEPSCs and mIPSCs, we examined how they are modulated by neuromodulators. Specifically, following the lead from a recent study on glutamate/GABA co-releases in the lateral habenula (17), we examined how adenosine and serotonin affect the minis in the striatum. We found that adenosine decreased the frequency but not the amplitude of both mEPSCs and mIPSCs (Fig. 3a–3d). The frequencies of both types of biphasic minis were also decreased (Fig. 3e and 3f). However, when compared to the chance of coincided mEPSCs and mIPSCs, which is related to the multiplicative effect of both mEPSC and mIPSC frequencies, the relative probability of both types of biphasic minis in fact increased (Fig. 3g and 3h). In contrast to adenosine, serotonin application did not alter the mini frequencies (Supplementary Fig. 1a–2h). Removal of external calcium also did not alter mini frequencies, indicating that the biphasic minis are not calcium triggered multivesicular release events (Supplementary Fig. 1i–2p). Overall, these results strengthen the notion that biphasic minis are unlikely the result of coincidently arrived mEPSCs and mIPSCs.

**Fig. 3.**
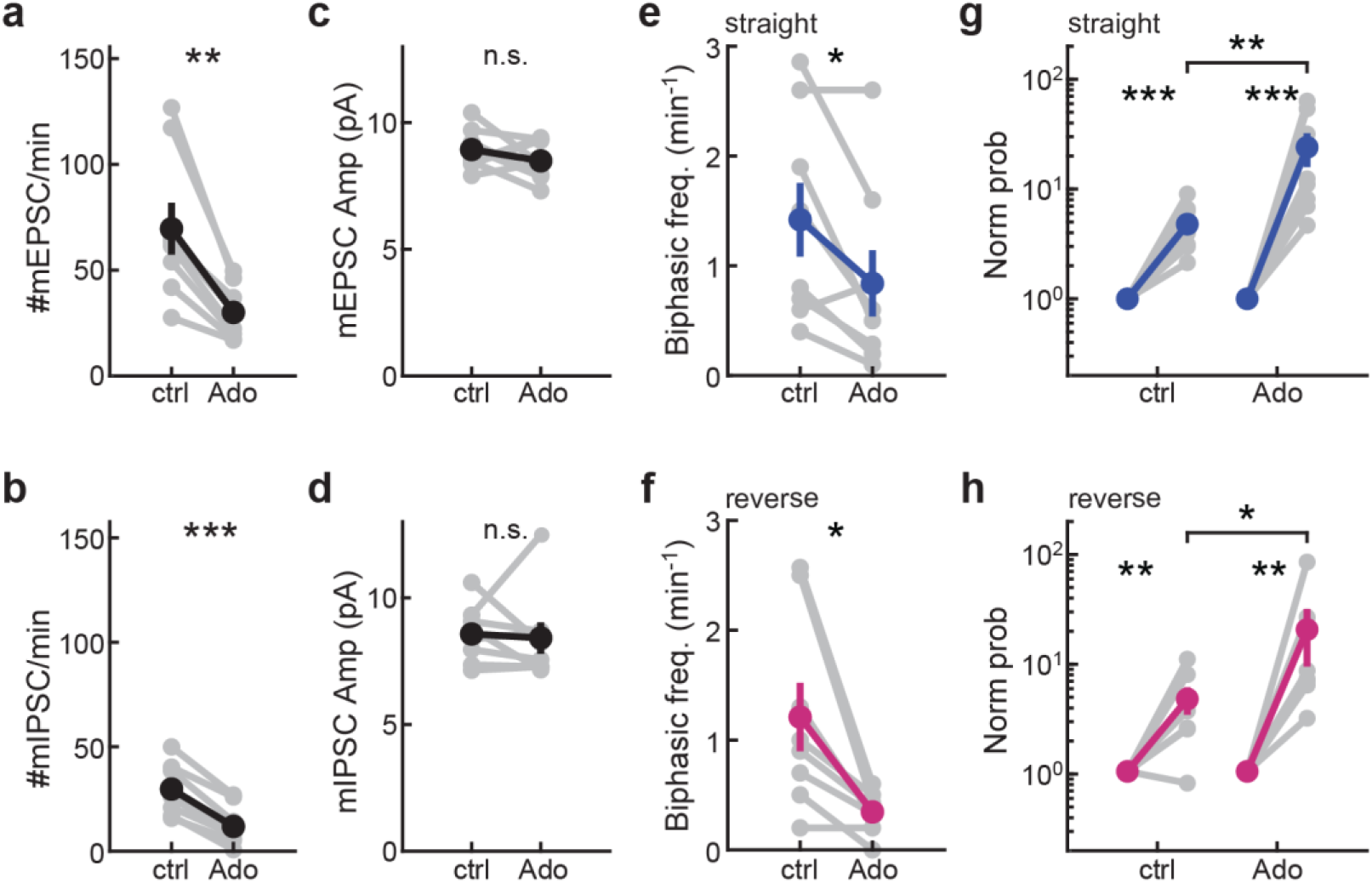
Adenosine decreased the number but not the probability of biphasic minis. **a**– **d**, Frequency of mEPSCs (**a**) and mIPSCs (**b**), and amplitudes of mEPSCs (**c**) and mIPSCs (**d**) of SPN neurons before (ctrl) and after the bath application of 20 μM adenosine (Ado). **e** and **f**, Frequency of straight (**e**) and reverse (**f**) biphasic minis before (ctrl) and after adenosine application. **g** and **h**, The probability of observed straight (**g**) or reverse biphasic minis (**h**) normalized to the probability of their occurrence due to coincidence of independent mEPSC and mIPSC in the same cells before (ctrl) and after adenosine application. All recordings were done at -30 mV. n (neurons/slices/mice) = 8/8/3 for both control and adenosine application.

### Dopaminergic axons are not the major source of biphasic minis

Because dopaminergic axons can release both glutamate and GABA (7, 26), we asked whether they might contribute to the biphasic minis. First, we performed minimal optogenetic stimulations of dopamine axon terminals. A single short pulse of blue light (∼470 nm, 1 ms, ∼2.5 mW/mm^2^) was applied to a field (Φ = 440 μm) centered on the recorded SPN in slices prepared from *DAT-Cre/Ai32* double heterozygous mice, which express channelrhodopsin (ChR2[H134R]-EYFP) in dopaminergic axons. The light intensity was adjusted so that failure and apparent quantal synaptic responses could be observed in different trials (Fig. 4a). Four types of responses were observed: failures, EPSCs only, IPSCs only, and biphasic PSCs (Fig. 4a and 4b). However, the probability of biphasic PSCs was not statistically higher than chance in this experiment (p = 0.29, sign test; Fig. 4c).

**Fig. 4.**
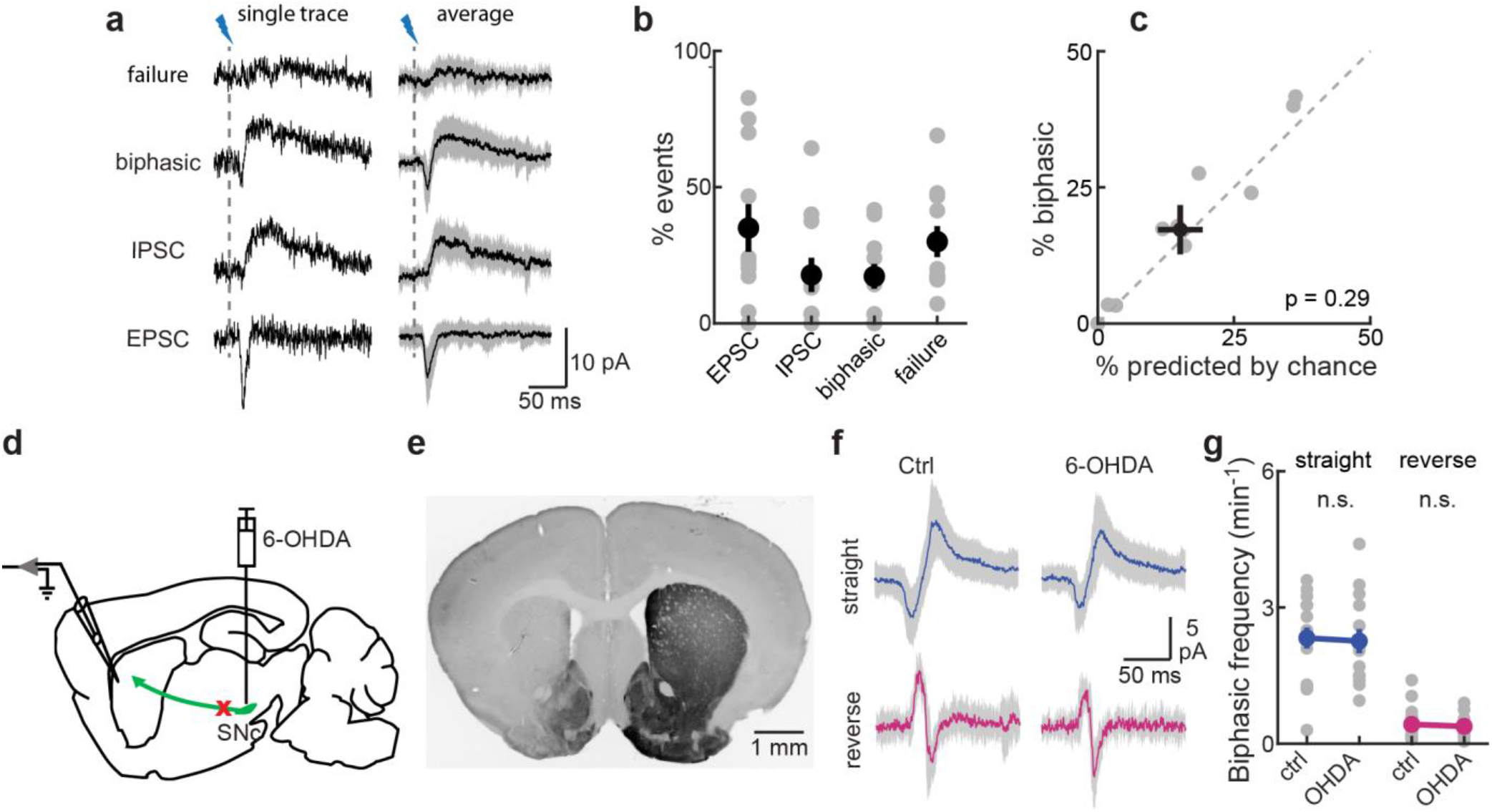
Dopaminergic axons are not the major source of biphasic minis. **a**, Example single events observed upon minimal optogenetic stimulation with the same cell (left) and the class average (right). **b**, Percentage of different type of events. n (neurons/slices/mice) = 11/8/3. **c**, Scatter plots of chance versus probability of measured biphasic event frequency. **d** and **e**, Unilateral injection of 6-OHDA assessed by tyrosine hydroxylase (TH) immunofluorescence in the striatum after 4 days. **f** and **g**, Example traces of straight and reverse biphasic minis (**f**) and their frequency (**g**) recorded in control and 6-OHDA lesion sides. n (neurons/slices/mice) = 16/8/4 for ctrl, and 14/7/4 for 6-OHDA. All recordings were done at -30 mV.

Second, we unilaterally injected 6-hydroxydopamine (6-OHDA) into SNc to deplete dopaminergic neurons (Fig. 4d). Near complete removal of tyrosine hydroxylase-positive axon terminals in the ipsilateral dorsal striatum was observed after 4 days (Fig. 4e). At day 4 and 5 post injection, comparable frequencies of both straight and reverse biphasic minis were observed between the control and the injected hemispheres (Fig. 4f and 4g; p= 0.68 for straight and 0.83 for reverse, Wilcoxon rank sum test). There were also no differences in frequency and amplitude of monophasic mEPSCs and mIPSCs (Supplementary Fig. 1). Together, these results suggest that dopamine axon terminals are not the main source of either biphasic or monophasic minis in the dorsolateral striatum.

### Biphasic minis are prevalent throughout the brain

Because our earlier results indicated that biphasic minis are prevalent across neuronal types in the striatum, we asked whether they are also prevalent across brain regions. In addition to SPNs and ChINs in the dorsolateral striatum, we recorded in the following brain regions of wildtype mice (Fig. 5a and Supplementary Fig. 1): barrel cortex (BC), basolateral amygdala (BLA), CA1 region of hippocampus (CA1), cerebellum (Cb), globus pallidus externa (GPe), medial prefrontal cortex (mPFC), suprachiasmatic nucleus (SCN), substantia nigra pars compacta (SNc), ventroposterior thalamus (Th) and primary visual cortex (V1). Surprisingly, we observed biphasic minis in all regions.

**Fig. 5.**
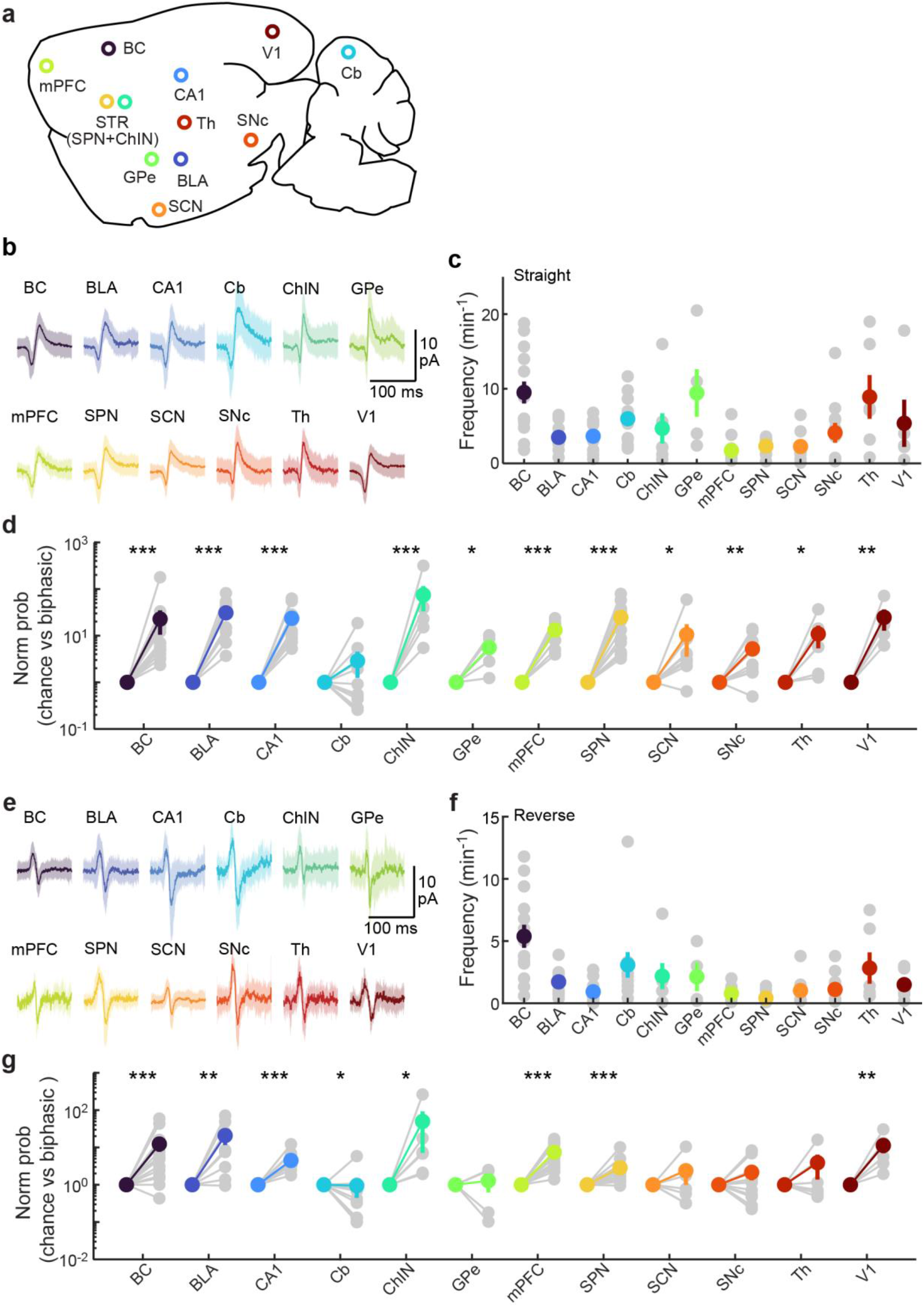
Biphasic minis are prevalent throughout the brain. **a**, Brain regions where electrophysiological recordings were done: the barrel cortex (BC), basolateral amygdala (BLA), CA1 region of hippocampus (CA1), cerebellum (Cb), cholinergic interneurons from dorsolateral striatum (ChIN), globus pallidus externa (GPe), medial prefrontal cortex (mPFC), spiny projection neurons from dorsolateral striatum (SPN), suprachiasmatic nucleus (SCN), substantia nigra pars compacta (SNc), ventral thalamus (Th), primary visual cortex (V1). **b**–**d**, Example of average traces (**b**), frequency (**c**), and probability normalized to chance (**d**) of straight biphasic minis across brain regions and cell types. **e**–**g**, Example of average traces (**e**), frequency (**f**), and probability normalized to chance (**g**) of straight biphasic minis across brain regions and cell types. From left to right in panels **c**, **d**, **f** and **g**, n (neurons/slices/mice) = 14/6/3, 9/7/3, 10/4/2, 11/6/3, 7/3/2, 5/4/2, 11/3/3, 16/7/4, 8/3/3, 11/6/3, 6/4/3, and 5/3/2. All recordings were done at ∼ -30 mV.

Straight biphasic minis occurred at a frequency ∼1–20 events/min that varied across brain regions (Fig. 5b and 5c). When compared with their chance levels calculated from measured mEPSC and mIPSC frequencies (Supplementary Fig. 1), straight biphasic minis occurred above chance in all regions except the cerebellum (Fig. 5d). Reverse biphasic minis had a frequency ∼1–12 events/min (Fig. 5e and 5f) and was above chance in seven out of 12 brain regions/cell types (Fig. 5g). In the barrel cortex, we also verified that both types of biphasic minis could be abolished by either NBQX or GABAzine (Supplementary Fig. 1). These results suggest that glutamate/GABA co-release is widespread across brain regions.

## DISCUSSION

Here, we investigated GABA and glutamate co-release using electrophysiological recordings of miniature postsynaptic currents. We found that biphasic minis involved both AMPA and GABA_A_ receptors were prevalent throughout many brain regions. Statistical analysis showed that the observed biphasic minis cannot be explained by coincided independent release of GABA and glutamate from independent vesicles.

Although many studies have shown evidence of axons co-transmitting glutamate and GABA in the brain (6–16, 23, 30, 36, 39–45), evidence for glutamate and GABA co-release is scarce with the exception of those from Entopeduncular nucleus (EP) axons onto the lateral habenula (9, 17). We found that eleven out of the 12 tested brain regions or cell types exhibited glutamate/GABA biphasic minis, which are thought to likely represent the co-release of these two opposing transmitters from a single vesicle (3, 4, 9, 36–38, 46). While surprising, this finding is consistent with recent studies showing that a small percentage of synaptic vesicles co-express both glutamate and GABA vesicular transporters in many brain regions (15, 18–21, 47). At the postsynaptic side, GABA_A_ and AMPA receptors have been found to colocalize at mossy fiber synapses (48).

Interestingly, we found two types of biphasic minis: ones in which the mEPSC component leading the mIPSC component (straight biphasic minis) and others with the mIPSC preceding the mEPSC (reverse biphasic minis). To our knowledge, the reverse biphasic mini has not been previously described. We verified that both type of biphasic minis depend on both AMPA and GABA_A_ receptors. Assuming that these biphasic minis arise from single-vesicular releases, the two types of biphasic minis likely reflect different synaptic organizations. For example, at some synapses giving rise to straight biphasic minis, AMPA receptors are more central to the active zone than GABA_A_ receptors, whereas at synapses with reverse biphasic minis, GABA_A_ receptors are closer to the release site. While it has been shown that AMPAR and GABA_A_R may exist in the same synapses or spines (48–50), direct testing of this hypothesis will likely require EM tomography study of neuronal synapses. Finally, our data cannot rule out the possibility that glutamate and GABA are packaged into separate vesicles, but the release of these distinct vesicles are tightly coupled within milliseconds. Coupled multivesicular release within an active zone triggered by putatively single calcium channel opening events or by local calcium sparks has been observed in specialized synapses (51, 52). Another possibility is that the neurotransmitter release from one terminal diffuse to an adjacent synapse to increase its release probability within milliseconds (53). However, under our conditions the biphasic minis are not sensitive to external calcium concentrations, making these possibilities less likely. Regardless of the precise mechanism, our results show that coordinated, time-locked co-release of glutamate and GABA is surprisingly common in the brain.

What is the function of glutamate/GABA co-release? First, although biphasic minis account for only a small fraction of total minis, they may be dominant in particular presynaptic neuronal types to shape their function. There have been several suggested possible mechanisms. Co-localization of GABA_A_ receptors with AMPA receptors could rapidly inhibit the excitatory response, providing a fast and more targeted form of ‘surround’ inhibition (54). Both computer simulations and experimental evidence suggest that inhibition within a few milliseconds and in close proximity to glutamate receptor excitations are highly efficient in attenuating postsynaptic calcium response amplitudes without impacting voltage changes (49, 50, 55). The spatial specificity of calcium dynamics may also be enhanced (55). Additionally, VGLUT expressed in axon terminals may result in enhanced uptake of GABA into synaptic vesicles (i.e., vesicular synergy) (19, 37), thereby increasing GABA release. Finally, there may be potential implications in plasticity. Activation of presynaptic metabotropic glutamate receptors or GABA_B_ receptors could modulate local synaptic transmissions (3, 15, 27, 28, 56). Calcium influx through NMDARs have also been found to selectively potentiates inhibition from a subset of inhibitory synapses in somatostatin positive interneuron (57).

The cellular origin of glutamate/GABA co-release remains unknown. In the striatum, our result suggests that dopaminergic axons are not the major contributor to glutamate and GABA co-release. This result is consistent with recent investigations. There appears to be minimal colocalization between VGLUT2 and VMAT2, which is responsible for vesicular GABA transport in these axons (8, 42, 58), in dopaminergic axonal terminals in the striatum [(31–33, 59); but see also (35)]. Additionally, glutamate and dopamine releases exhibit different properties (31, 40, 59, 60). One possible source might be GABAergic interneurons axon terminals that also express VGLUT2 or VGLUT3. For instance, GABA neurons of the hypothalamic anteroventral periventricular nucleus express VGLUT2 (61). Likewise multiple populations of GABAergic interneurons in the cerebral cortex and hippocampus (43, 62, 63) and immature GABAergic synapses from neurons of the nucleus of the trapezoid body in the lateral superior olive (LSO) (44) express VGLUT3. Overall, it appears that neurons that release glutamate and GABA express either VGLUT2 or VGLUT3, but not VGLUT1 (45). Future revelation of the precise cellular source of these biphasic minis will enable targeted manipulation to dissect the function and plasticity of this wide-spread co-release of glutamate and GABA.

## METHODS

### Mice

Animal handling and experimental protocols were performed in accordance with the recommendations in the Guide for the Care and Use of Laboratory Animals, written by the National Research Council (US) Institute for Laboratory Animal Research, and were approved by the Institutional Animal Care and Use Committee (IACUC) of the Oregon Health & Science University (#IP00002274). *DAT-IRES-cre* (B6.SJL-*Slc6a3^tm1.1(cre)Bkmn^*/J; Jax #006660) homozygous and *Ai32* (B6;129S-*Gt(ROSA)26Sor^tm32(CAG-COP4*H134R/EYFP)Hze^*/J; Jax #012569) homozygous mice were bred for minimal optogenetic stimulation experiments. *D1-Tdtomato* (Jax #016204) hemizygous were bred with C57BL/6 mice. Both sexes of mice ∼ P45 (mean ± s.e.m. = 45 ± 2.7; range between P14–P114).

### Acute slice preparation

Mice were transcardially perfused with ice-cold, gassed aCSF containing (in mM) 127 NaCl, 25 NaHCO_3_, 10 D-glucose, 2.5 KCl, 1.25 NaH_2_PO_4_, 2 CaCl_2_, and 1 MgCl_2_. The brain was then resected and 300 µm-thick slices were obtained using a vibratome (Leica VT1200s) in an ice-cold, gassed sucrose-cutting solution containing (in mM): 210 sucrose, 25 NaHCO_3_, 10 D-glucose, 2.5 KCl, 7 MgCl_2_, 0.5 CaCl_2_, 1.25 NaH_2_PO_4_ and 1 sodium pyruvate. The slices were then incubated in gassed aCSF at 35°C for 30 minutes and subsequently kept at room temperature for up to 6 hours.

Coronal slices were made for the following brain regions: the dorsolateral striatum, barrel cortex (BC), basolateral amygdala (BLA), globus pallidus externa (GPe), medial prefrontal cortex (mPFC), suprachiasmatic nucleus (SCN), substantia nigra pars compacta (SNc), ventral thalamus (Th) and primary visual cortex (V1). Slices containing CA1 region of the hippocampus (CA1) were cut horizontally, and the cerebellum (Cb) were cut sagittally.

### Patch-clamp electrophysiology

Whole-cell voltage-clamped recordings were performed using a MultiClamp 700B amplifier (Molecular Devices) controlled with custom software written in MATLAB. Electrophysiological signals were filtered at 2 kHz before being digitized at 20 kHz. Slices were perfused at room temperature with gassed aCSF containing CPP (10 µM). Recording pipettes (3–5 MΩ), were pulled from borosilicate glass (G150F-3; Warner Instruments) using a model P-1000 puller (Sutter Instruments). Series resistance was 10–25 MΩ. The internal solution contained (in mM): 126 Cs-gluconate, 10 HEPES, 5 Na-phosphocreatine, 0.5 Na-GTP, 4 Na-ATP, 5 TEA-Cl, 5 EGTA and 4 QX-314 bromide with an osmolarity of 280-290 mOsmol/kg and pH ∼7.2 adjusted with CsOH. The junction potential was calculated to be -14 mV, as calculated using JCal from Clampex software (Molecular Devices). Voltages were not corrected for the theoretical liquid junction potentials. For IV curves recordings, we decreased the chloride concentration in the internal solution (in mM): 126 Cs-gluconate, 10 HEPES, 8 Na-phosphocreatine, 0.3 Na-GTP, 4 Mg-ATP, 1 EGTA and 1 QX-314 chloride with an osmolarity of 280-290 mOsmol/kg and pH ∼7.3 adjusted with CsOH.

For miniature postsynaptic currents recordings, TTX (1 μM) and CPP (10 µM) was added to the bath to block action potentials and NMDA currents, respectively. 20-minute traces were recorded in voltage-clamp mode and mEPSCs and mIPSCs events were detected using a template matching built-in feature in Clampfit (Molecular devices). Recordings were done in acute slices from C57BL/6 mice. We also did miniature recordings in SPN from the dorsolateral striatum from *D1-Tdtomato* mice. dSPNs were identified by the presence of fluorescence, meanwhile iSPNs were non-fluorescent neurons that was not ChINs, which can be identified via its unique, large soma.

Biphasic minis were visually identified via its unique shape. We only included biphasic minis in which the mEPSC peak was connected to the mIPSC peak by a smooth continuous line as have been observed elsewhere (9, 17). In other words, we excluded biphasic events in which, for straight biphasic minis, the decay of the mEPSC to the rise of the mIPSC and, for reverse biphasic minis, the decay of the mIPSC to the rise of the mEPSC were discontinuous. In pharmacological experiments, 10 μM gabazine and 10 μM NBQX were added to the aCSF to block GABA_A_R and AMPAR, respectively. In neuromodulation experiments, adenosine (20 µM) or serotonin (10 µM) were added to the perfusion solution.

For recording in different region and cell types, identification of the neurons was based mostly on their characteristic morphology and location. For each region, cells were held at a holding potential that maximize the detection of biphasic minis (V_H_ in mV): BC, - 25; BLA -30; CA1 -25; Cb -35; ChIN -25; GPe -40; mPFC -30; SCN -30; SNc -30; Th -30; and V1 -30. For recordings in Purkinje cells, 100 µM CdCl_2_ was added to the bath to avoid calcium spikes and calcium oscillations.

### 6-OHDA injections

6-hydroxydopamine (Tocris) injections were performed in WT mice at 8-9 weeks of age. Animals were anesthetized using 2 % isoflurane, mixed with oxygen, and placed in a stereotaxic frame. After making a small incision to expose the scalp, the skull was clean above bregma and lambda and a dental drill was used to make a small craniotomy above the Substantia Nigra pars compacta. 6-OHDA was dissolved in saline (0.9% w/v NaCl with 0.02% w/v ascorbic acid), to a final concentration of 5 μg/μl, immediately before use to minimize the oxidative effects on 6-OHDA. Injections were performed in 3 sites per animal (600 nl/site), at a rate ∼35 nl/min into the SNc (at AP 2.4 mm and ML 1.25 mm from bregma, and DV 3.8, 4 and 4.2 mm, respectively). Injections were performed using a micropipette pulled using a micropipette puller (Sutter Instrument, Co.). Desipramine (25 mg/kg delivered IP) was given 30 min prior to 6-OHDA infusion to block uptake of the toxin by noradrenergic neurons.

4 days after unilateral 6-OHDA injection, electrophysiological recordings were done in both sides. Dopaminergic lesion was confirmed by immunohistochemistry of striatal tyrosine hydroxylase (TH) which showed complete depletion of TH in the striatal hemisphere ipsilateral to the 6-OHDA lesion.

### Estimation of onset delay by fitting

The biphasic minis were fitted using the averaged mEPSC and mIPSC trace (those in Supplementary Fig. 1A, -30 mV) with freedom in amplitudes and the relative onset delays. The fitting was carried out in MATLAB using custom-made algorithms based on nonlinear least-squares data fitting by the Gauss-Newton method with the Levenberg-Marquardt adjustment. Near all traces (181 out of 184 straight biphasic minis and all 151 reverse biphasic minis) were well fitted by this approach. The averaged onset delay was ∼10 ms for both biphasic minis types. We therefore used this value as the time window to calculate chance probability (below).

### Probabilistic analysis of biphasic minis

To determine whether the observed number of biphasic minis events was due to chance, we calculated the expected probability of two independent events (one mEPSC and one mIPSC) to occur in the same time window (10 ms). The independent probability (i.e. chance) was calculated as the joint probability of finding an mEPSC (pE) or mIPSC (pI) in a 10 ms time window:

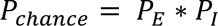

Where P_E_ corresponds to the chance of mEPSCs found in 10 ms and P_I_ corresponds to the chance of mIPSCs found in 10 ms.

The independent probability was compared to P(chance), i.e., the probability of finding a biphasic mini in a 10 ms time window. Where P(chance) corresponds to the # of biphasic minis found in 10 ms. In the analysis, P(chance) was normalized by the independent probability.

### Optogenetic

Recordings were done from coronal slices (300 µm thickness) containing the dorsolateral striatum from *DAT-Cre*/*Ai32* double-het mice, which expresses ChR2(H134R)-EYFP in dopaminergic axons. Minimal optogenetic stimulation of dopaminergic axons was evoked by applying a single short pulse (1 ms) of blue light (LED) under widefield illumination under the 60x objective at an empirically determined low intensity light. Light was applied to a field (440 μm) centered on the recorded neuron, and the intensity was adjusted so that failure and apparent quantal synaptic responses could be observed in different trials. In the voltage-clamp mode, cells were held at a holding potential (V_H_) of -45 mV to record biphasic responses (EPSCs and IPSCs). At this V_H_, mEPSCs and mIPSCs and upward and downward responses of biphasic currents had similar amplitude. Pulses were applied every 30 s and 30 trials were recorded.

For minimal stimulation experiments, the expected percentage of biphasic PSCs generated by chance from independent EPSCs and IPSCs were calculated from the following equation:

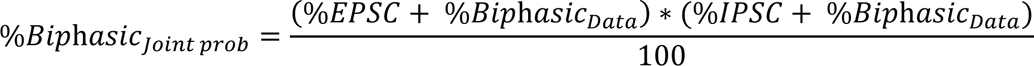

### Data analysis, presentation, and statistics

Quantification and statistical tests were performed using custom software written in MATLAB. Paired and unpaired t-tests were used for paired and unpaired data, respectively, unless otherwise noted. Averaged data are presented as mean ± SEM, unless noted otherwise. Throughout the paper, “n” indicates the number of neurons. In all figures, *: p ≤ 0.05 and is statistically significant, **: p ≤ 0.01, and ***: p ≤ 0.001.

## ACKNOWLEDGEMENTS

We thank Drs. Paul Brehm, Gary Westbrook, John Williams, and James Jones for critical comments on the manuscript. We thank all members of the Mao and Zhong laboratories at the Vollum Institute for constructive discussions. This work was supported by two NIH BRAIN Initiative awards (RF1MH130784 and R01NS104944) and an NINDS R01 (R01NS127013) to H.Z.

## AUTHOR CONTRIBUTIONS

C.C. made the initial observation of biphasic minis. C.C. and H.Z. designed the experiments. C.C performed the experiments and data analyses. L.M performed 6-OHDA injections. M.Q. maintained mouse husbandry. H.Z. and C.C wrote the manuscript. All authors edited and commented on the manuscript. H.Z. secured the funding and supervised the project.

## FIGURES AND FIGURE LEGENDS

**Supplementary Fig. 1.**
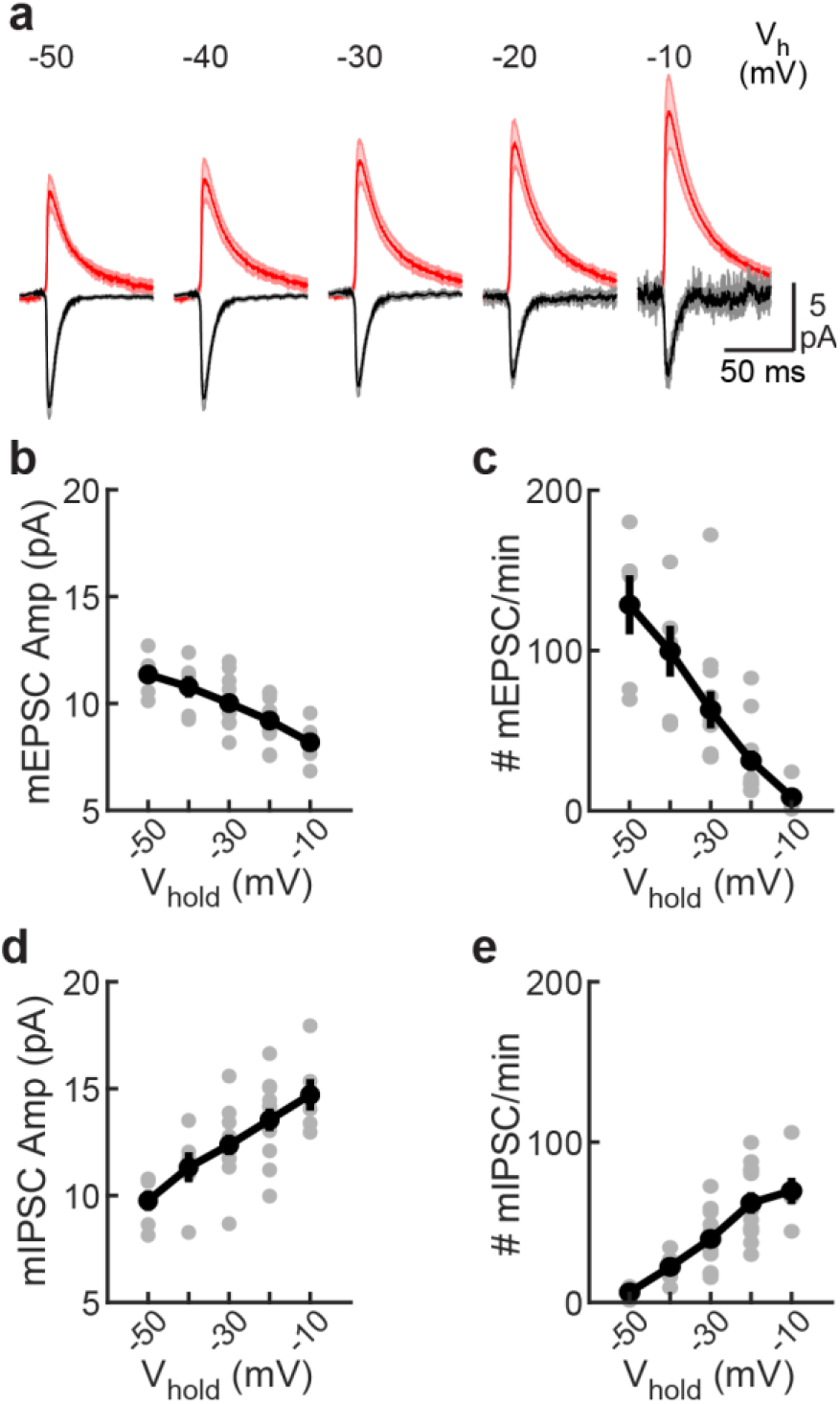
Voltage dependence of monophasic minis. **a**, Average traces per V_H_ (-50, -40, -30, -20 and -10 mV) of monophasic mEPSC and mIPSC recorded in SPN dlSTR. Monophasic mEPSC and mIPSC amplitudes behave as expected by changes in driving force. **b**, Voltage-dependent amplitude of mEPSC. **c**, Voltage-dependent frequency of mEPSC. **d**, Voltage-dependent amplitude of mIPSC. **e**, Voltage-dependent frequency of mIPSC. Data are from 5–11 neurons in 7 slices and 4 mice.

**Supplementary Fig. 2.**
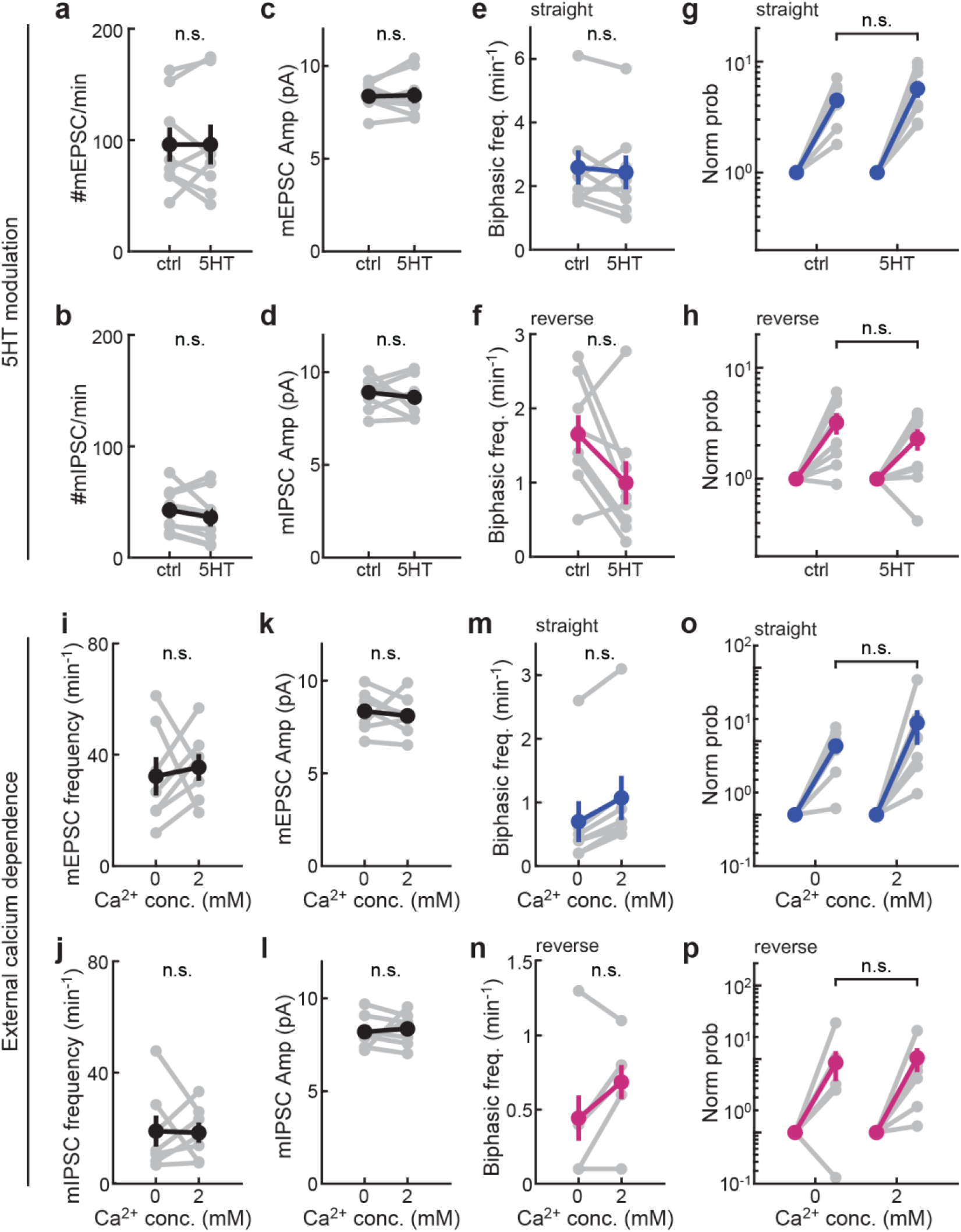
Serotonin and calcium exhibited no significant effect on biphasic minis frequencies. **a**–**d**, Frequency of mEPSCs (**a**) and mIPSCs (**b**), and amplitudes of mEPSCs (**c**) and mIPSCs (**d**) of SPN neurons before (ctrl) and after the bath application of 10 μM serotonin (5HT). **e** and **f**, Frequency of straight (**e**) and reverse (**f**) biphasic minis before (ctrl) and after serotonin application. **g** and **h**, The probability of observed straight (**g**) or reverse biphasic minis (**h**) normalized to the probability of their occurrence due to coincidence of independent mEPSC and mIPSC in the same cells before (ctrl) and after serotonin application. n (neurons/slices/mice) = 8/8/3 for both control and serotonin application. **i**– **l**, Frequency of mEPSCs (**i**) and mIPSCs (**j**), and amplitudes of mEPSCs (**k**) and mIPSCs (**l**) of SPN neurons under normal (2 mM) and decreased (0 mM) calcium concentrations. **m** and **n**, Frequency of straight (**m**) and reverse (**n**) biphasic minis in the presence of 0 or 2 mM extracellular calcium concentrations. **o** and **p**, The probability of observed straight (**o**) or reverse biphasic minis (**p**) normalized to the probability of their occurrence due to coincidence of independent mEPSC and mIPSC in the presence of 0 or 2 mM extracellular calcium. n (neurons/slices/mice) = 7/7/2 for both conditions. All recordings were done at - 30 mV.

**Supplementary Fig. 3.**
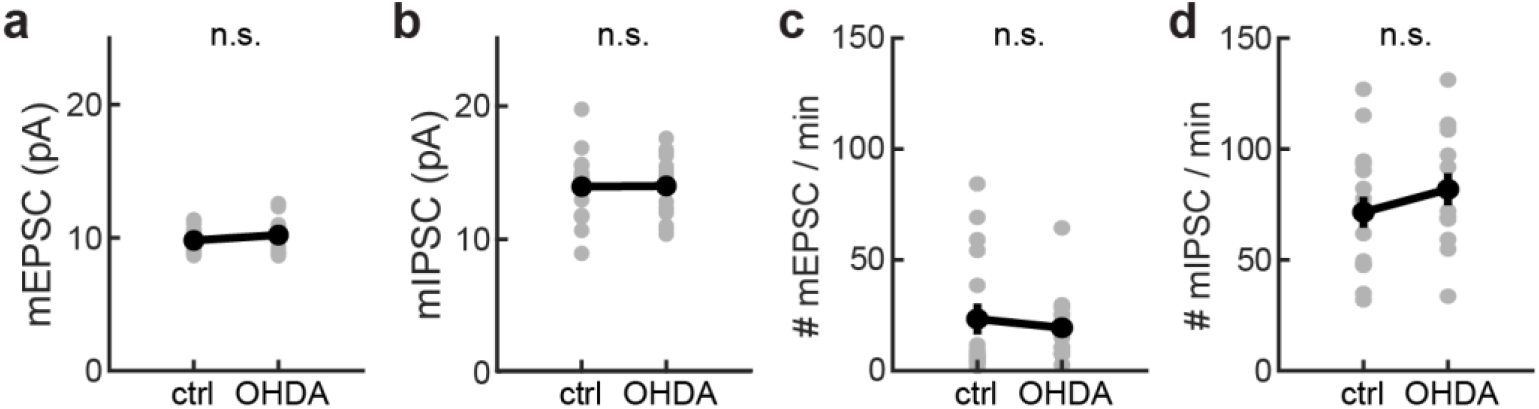
6-OHDA effects on monophasic minis. **a**, mEPSC amplitude of control vs 6-OHDA lesion side from dlSTR SPN. **b**, mIPSC amplitude of control vs 6-OHDA lesion side. **c**, mEPSC frequency of control vs 6-OHDA lesion side. **d**, mIPSC frequency of control vs 6-OHDA lesion side. n (neurons/slices/mice) = 16/8/4 for ctrl and 14/7/4 for OHDA. All recordings were done at -30 mV.

**Supplementary Fig. 4.**
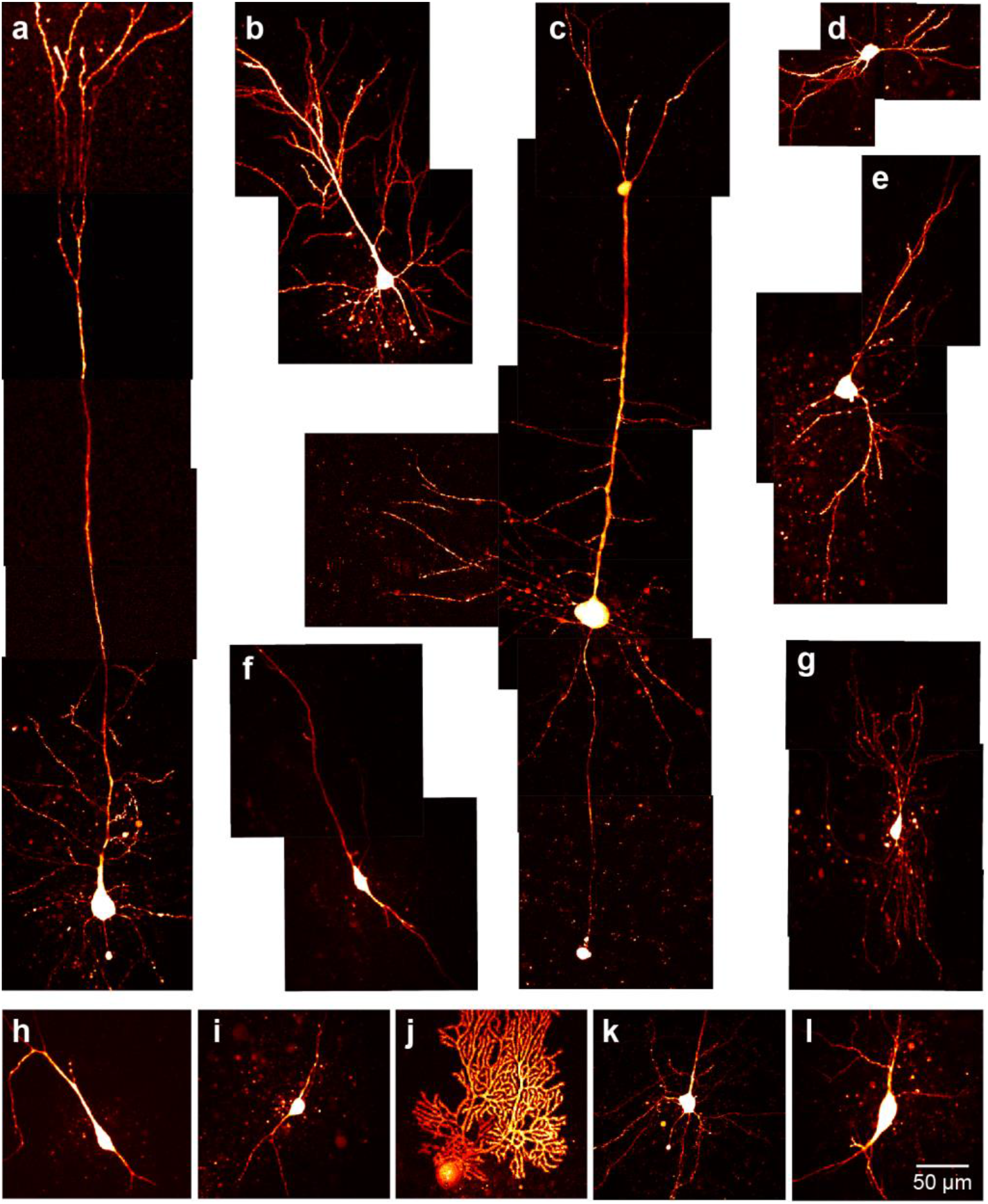
2P z-stack of morphological reconstruction of neurons. Example of neurons in which patch-clamp recordings were performed. **a**, Layer IV/V pyramidal neuron (BC). **b**, CA1 pyramidal neurons (CA1). **c**, Layer IV/V pyramidal neuron (V1). **d**, Spiny projection neuron (dlSTR). **e**, Pyramidal neurons (BLA). **f**, GABAergic neuron (GPe). **g**, Thalamic neuron (Th). **h**, Dopaminergic neuron (SNc). **i**, Gabaergic neuron (SCN). **j**, Purkinje cell, Lobule VI (Cb). **k**, Layer VI pyramidal neurons (mPFC). **l**, Cholinergic interneurons (dlSTR).

**Supplementary Fig. 5.**
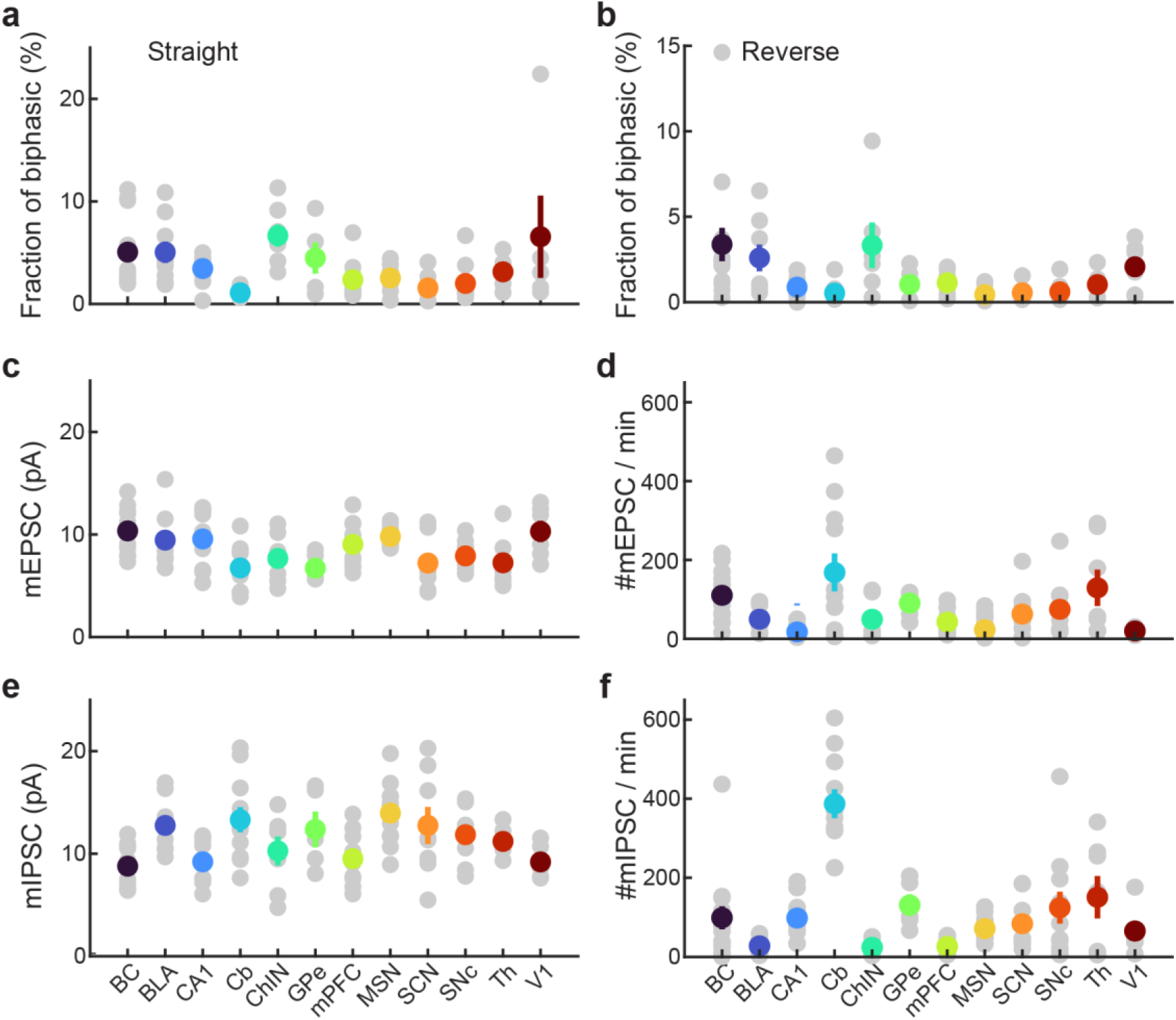
Parameters of monophasic minis across brain regions or neuronal types. **a**, Percentage of minis that correspond to straight biphasic minis compared to the total of monophasic minis (mEPSC + mIPSC) for all brain regions. **b**, Same as in panel **a** but for reverse biphasic minis. **c**, mEPSC amplitude for all brain regions. **d**, mEPSC frequency for all brain regions. **e**, mIPSC amplitude for all brain regions. **f**, mIPSC frequency for all brain regions. Abbreviations: Barrel cortex (BC), Basolateral Amygdala (BLA), CA1 region of Hippocampus (CA1), Cerebellum (Cb), Cholinergic interneurons from dorsolateral striatum (ChIN), Globus pallidus externa (GPe), medial prefrontal cortex (mPFC), Spiny projection neurons from dorsolateral striatum (SPN), Suprachiasmatic nucleus (SCN), Substantia nigra pars compacta (SNc), Ventral thalamus (Th) and Primary visual cortex (V1). From left to right in each panel, n (neurons/slices/mice) = 14/6/3, 9/7/3, 10/4/2, 11/6/3, 7/3/2, 5/4/2, 11/3/3, 16/7/4, 8/3/3, 11/6/3, 6/4/3, and 5/3/2.

**Supplementary Fig. 6.**
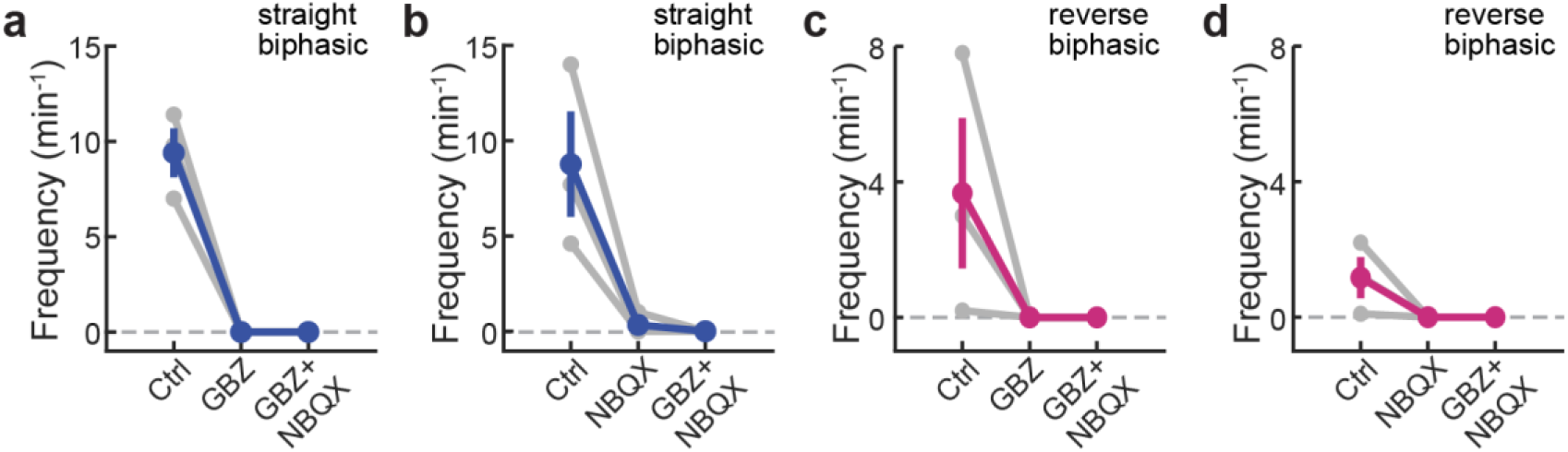
Both types of biphasic minis in the barrel cortex are blocked by either GABAAR or AMPAR antagonist. **a**, GABAAR antagonist (Gabazine, GBZ) abolished straight biphasic minis in Layer IV/V pyramidal neurons from the barrel cortex. **b**, AMPAR antagonist (NBQX) abolished straight biphasic minis. **c**, Gabazine abolished reverse biphasic minis. **d**, NBQX abolished reverse biphasic minis. n (neurons/slices/mice) = 3/3/2. All recordings were done at ∼ -30 mV.

